# Binaural summation of amplitude modulation involves weak interaural suppression

**DOI:** 10.1101/278192

**Authors:** D.H. Baker, G. Vilidaite, E. McClarnon, E. Valkova, A. Bruno, R.E. Millman

**Affiliations:** Department of Psychology, University of York, Heslington, York, YO10 5DD, UK; York Biomedical Research Institute, University of York, Heslington, York, YO10 5DD, UK; School of Psychology, University of Southampton, University Road, Southampton, SO17 1BJ, UK; Manchester Centre for Audiology and Deafness, University of Manchester, Manchester Academic Health Science Centre, M13 9PL, UK; NIHR Manchester Biomedical Research Centre, Central Manchester University Hospitals NHS Foundation Trust, Manchester Academic Health Science Centre, Manchester, M13 9WL, UK

**Keywords:** binaural combination, EEG, amplitude modulation

## Abstract

The brain combines sounds from the two ears, but what is the algorithm used to achieve this summation of signals? Here we combine psychophysical amplitude modulation discrimination and steady-state electroencephalography (EEG) data to investigate the architecture of binaural combination for amplitude-modulated tones. Discrimination thresholds followed a ‘dipper’ shaped function of pedestal modulation depth, and were consistently lower for binaural than monaural presentation of modulated tones. The EEG responses were greater for binaural than monaural presentation of modulated tones, and when a masker was presented to one ear, it produced only weak suppression of the response to a signal presented to the other ear. Both data sets were well-fit by a computational model originally derived for visual signal combination, but with suppression between the two channels (ears) being much weaker than in binocular vision. We suggest that the distinct ecological constraints on vision and hearing can explain this difference, if it is assumed that the brain avoids over-representing sensory signals originating from a single object. These findings position our understanding of binaural summation in a broader context of work on sensory signal combination in the brain, and delineate the similarities and differences between vision and hearing.

## Introduction

The auditory system integrates information across the two ears. This operation confers several benefits, including increased sensitivity to low intensity sounds [1] and inferring location and motion direction of sound sources based on interaural time differences [2]. In some animals, such as bats and dolphins, echolocation can be precise enough to permit navigation through the environment, and there are reports of visually impaired humans using a similar strategy [3,4], which requires both ears [5]. But what precisely is the algorithm that governs the combination of sounds across the ears? The nonlinearities inherent in sensory processing mean that simple linear signal addition is unlikely. This study uses complementary techniques (psychophysics, steady-state electroencephalography (EEG) and computational modelling) to probe the neural operations that underpin binaural summation of amplitude-modulated signals.

Classical psychophysical studies demonstrated that the threshold for detecting a very faint tone is lower when the tone is presented binaurally versus monaurally. Shaw et al. [1] presented signals to the two ears that were equated for each ear’s individual threshold sound level when presented binaurally. This accounted for any differences in sensitivity (or audibility), and revealed that summation (the improvement in sensitivity afforded by binaural presentation) was approximately 3.6 dB (a factor of 1.5). Subsequent studies have provided similar or slightly lower values [i.e. 6–8], and there is general agreement that two ears are better than one at detection threshold [9]. This difference persists above threshold, with intensity discrimination performance being better binaurally than monaurally [10]. Furthermore, binaural sounds are perceived as being slightly louder than monaural sounds, though typically less than twice as loud [11–14].

When a carrier stimulus (typically either a pure-tone or broadband noise) is modulated in amplitude, neural oscillations at the modulation frequency can be detected at the scalp [15–19], being typically strongest at the vertex in EEG recordings [20]. This steady-state auditory evoked potential (SSAEP) is typically greatest around 40 Hz [20,21] and increases monotonically with increasing modulation depth [17,18]. For low signal modulation frequencies (<55 Hz), brain responses are thought to reflect cortical processes [15,20–22]. The SSAEP has been used to study binaural interactions, showing evidence of interaural suppression [23,24] and increased responses from binaurally fused stimuli [17,22].

The perception of amplitude-modulated stimuli shows similar properties to the perception of pure-tones in terms of binaural processing. For example, binaural sensitivity to amplitude modulation (AM) is better than monaural sensitivity [25,26], and the perceived modulation depth is approximately the average of the two monaural modulation depths over a wide range [27]. Presenting two different modulation frequencies to the left and right ears can produce the percept of a ‘binaural beat’ pattern at the difference intermodulation frequency (the highest minus the lowest frequency), suggesting that the two modulation frequencies are combined centrally [28]. Finally, both the detection of intensity increments and detection of AM [29] follows Weber-like behaviour [30,31] at higher pedestal levels (i.e. Weber fractions for discrimination are approximately constant with pedestal level) similar to that typically reported for intensity discrimination [32]. However, despite these observations, detailed investigation and modelling of the binaural processing of amplitude-modulated tones is lacking.

Computational predictions for both psychophysical and electrophysiological results can be obtained from parallel work that considers the combination of visual signals across the left and right eyes. In several previous studies, a single model of binocular combination has been shown to successfully account for the pattern of results from psychophysical contrast discrimination and matching tasks [33,34], as well as steady-state EEG experiments [35]. The model, shown schematically in Figure 1a, takes contrast signals (sinusoidal modulations of luminance) from the left and right eyes, which mutually inhibit each other before being summed as follows:

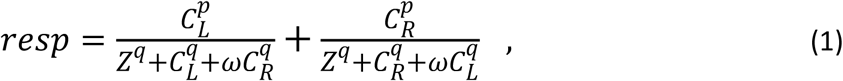

where *resp* is the model response, *C*_*L*_ and *C*_*R*_ are the contrast signals in the left and right eyes respectively, *ω* is the weight of interocular suppression, *Z* is a constant governing the gain of the model, and *p* and *q* are exponents with the typical constraint that *p*>*q*. In all experiments in which the two signals have the same visual properties [33–35], the weight of interocular suppression has a value around *ω* = 1 (and is assumed to be effectively instantaneous, though in reality is likely subject to some delay).

**Figure 1:**
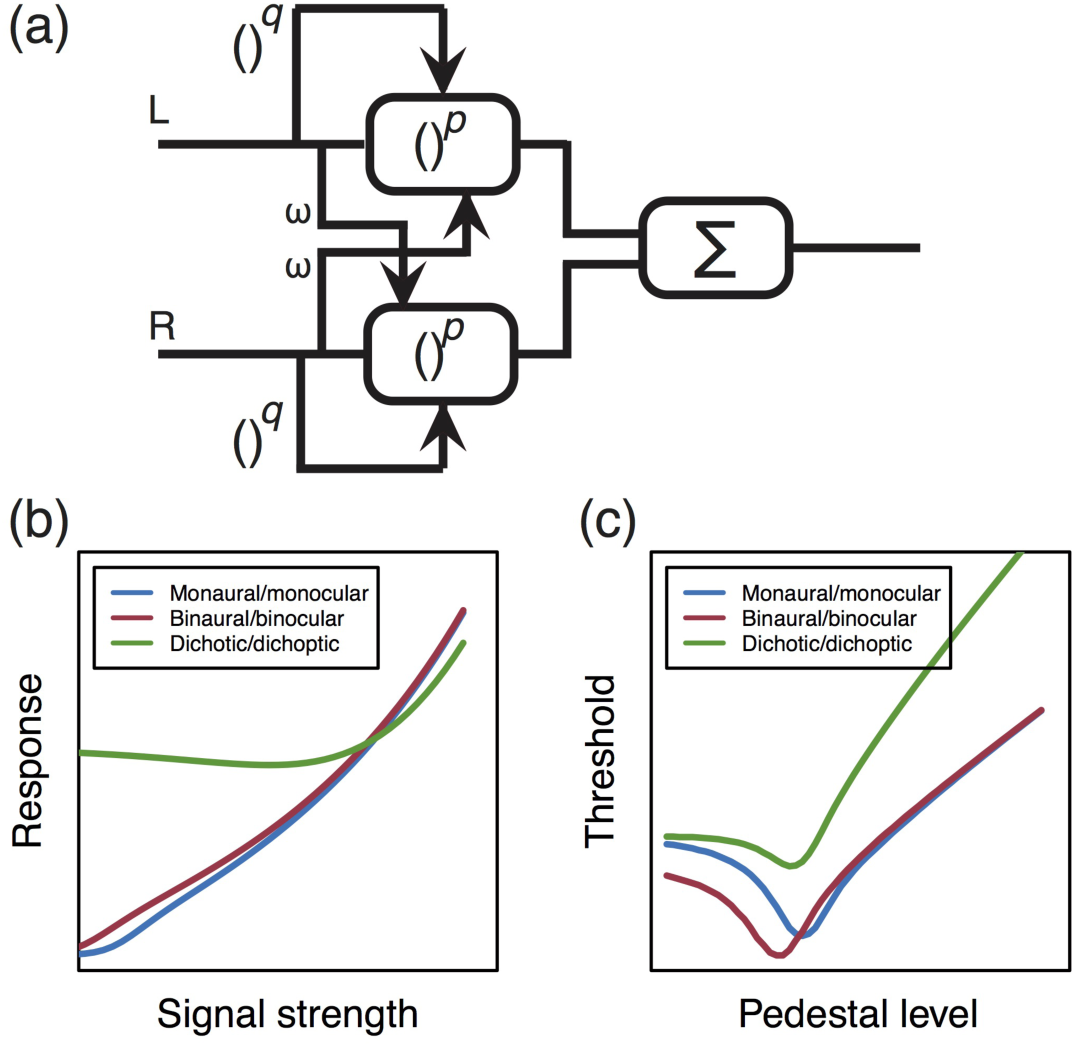
Schematic of signal combination model and qualitative predictions. Panel (a) shows a diagram of the signal combination model, which incorporates weighted inhibition between left and right channels before signal combination (Σ). Panel (b) shows the predictions of this model for various combinations of inputs to the left and right channels, as described in the text. Predictions for discrimination of increases in modulation depth for similar conditions are shown in panel (c).

Whereas vision studies typically modulate luminance relative to a mean background level (i.e. contrast), in hearing studies the amplitude modulation of a carrier waveform can be used to achieve the same effect. We can therefore test empirically whether binaural signal combination is governed by the same basic algorithm (in the tradition associated with David Marr [36]) as binocular signal combination by replacing the *C* terms in equation 1 with modulation depths for AM stimuli.

The response of the model for different combinations of inputs is shown in Figure 1b, with predictions being invariant to the sensory modality (hearing or vision). In the monaural/monocular (“mon”) condition (blue), signals are presented to one channel only. In the binaural/binocular (“bin”) condition (red) equal signals are presented to the two channels. In the dichotic/dichoptic (“dich”) condition (green) a signal is presented to one channel, with a fixed high amplitude ‘masker’ presented to the other channel throughout. For *ω* = 1, the *mon* and *bin* conditions produce similar outputs, despite a doubling of the input (two channels vs one). This occurs because the strong suppression between channels offsets the gain in the input signal. This pattern of responses is consistent with the amplitudes recorded from steady-state visual evoked potential experiments testing binocular combination in humans [35].

The model response can also be used to predict the results of psychophysical increment detection experiments in which thresholds are measured for discriminating changes in the level of a ‘pedestal’ stimulus (e.g. a stimulus of fixed intensity). In these experiments, thresholds are defined as the horizontal translation required to produce a unit increase vertically along the functions in Figure 1b. In other words, psychophysical performance measures the *gradient* of the contrast response function. These predictions are shown in Figure 1c and have a characteristic ‘dipper’ shape, in which thresholds first decrease (facilitation), before increasing (masking). The *mon* and *bin* functions converge at higher pedestal levels, and the *dich* function shows strong threshold elevation owing to the suppression between the two channels (when *ω* = 1). Again, this pattern of functions is consistent with those reported in psychophysical studies of binocular vision [34].

The present study uses two complementary methods – psychophysical AM depth discrimination, and steady-state auditory evoked potentials – to investigate binaural signal combination in the human brain. The results are compared with the predictions of the computational model [34,35] described above (see Figure 1) and modifications to the model are discussed in the context of functional constraints on the human auditory system. This principled, model-driven approach positions our understanding of binaural summation in a broader context of work on sensory signal combination in the brain.

## Methods

### Apparatus & stimuli

Auditory stimuli were presented over Sennheiser (HD 280 pro) headphones (Sennheiser electronic GmbH, Wedemark, Germany), and had an overall presentation level of 80 dB SPL. An AudioFile device (Cambridge Research Systems Ltd., Kent, UK) was used to generate the stimuli with a sample rate of 44100 Hz. Stimuli consisted of a 1-kHz pure-tone carrier, amplitude modulated at a modulation frequency of either 40 Hz or 35 Hz (see Figure 2), according to the equation:

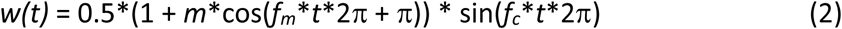

**Figure 2:**
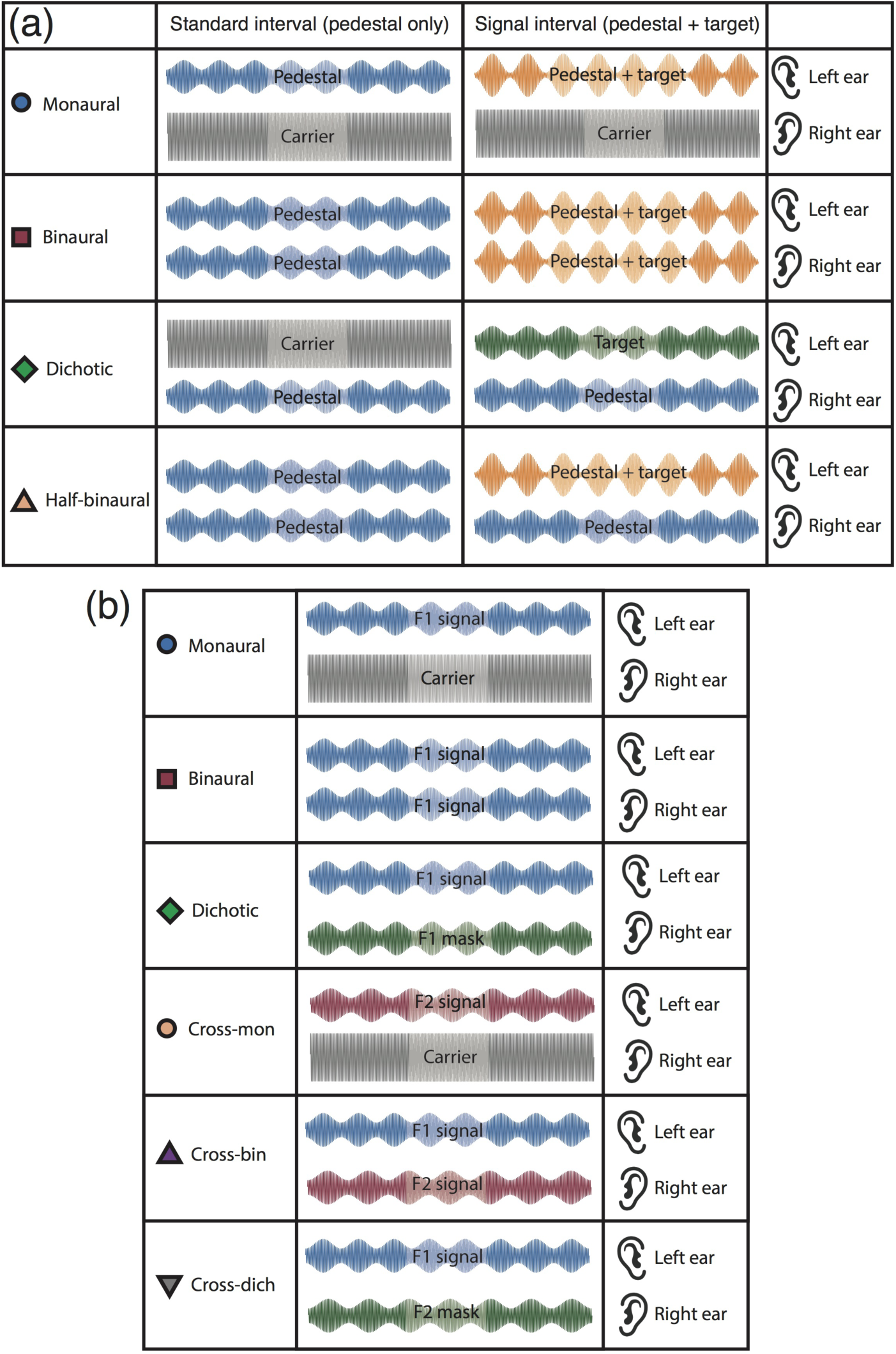
Summary of conditions and stimuli. Panel (a) illustrates the arrangement of pedestal and target modulations for the psychophysics experiment in the standard (pedestal only) interval (left) and signal (pedestal + target) interval (right) for four different interaural arrangements (rows). In all cases, the modulation frequency was 40 Hz and participants were asked to indicate the interval containing the target. A range of pedestal modulation depths were tested, with target modulation depths determined by a staircase algorithm. Panel (b) shows stimulus arrangements for six conditions in the EEG experiment. Stimuli designated ‘signal’ had different modulation depths in different conditions, whereas stimuli designated ‘masker’ had a fixed modulation depth (*m* = 50%) for all signal modulation depths. In all experiments stimulation was counterbalanced across the two ears, so the left ear/right ear assignments here are nominal.

where *f*_*m*_ is the modulation frequency in Hz, *f*_*c*_ is the carrier frequency in Hz, *t* is time in seconds, and *m* is the modulation depth, with a value from 0 - 1 (though hereafter expressed as a percentage, 100\**m*). We chose not to compensate for overall stimulus power (as is often done for AM stimuli, e.g. [37]) for 3 reasons (see also [29]). First, such compensation mostly affects AM detection thresholds at much higher modulation frequencies than we used here [e.g. see Figure A1 of 38]. Second, compensation makes implicit assumptions about the cues used by the participant in the experiment, and we prefer to make any such cues explicit through computational modelling. Third, we confirmed in a control experiment that compensation had no systematic effect on thresholds (see Supplementary Figure S1). The modulation depth and the assignments of modulation frequencies delivered to the left and right ears were varied parametrically across different conditions of the experiments.

EEG data were recorded with a sample frequency of 1 kHz using a 64-electrode Waveguard cap and an ANT Neuroscan (ANT Neuro, Netherlands) amplifier. Activity in each channel was referenced to the whole head average. Signals were digitised and stored on the hard drive of a PC for later offline analysis. Stimulus onset was coded on the EEG trace using digital triggers sent via a BNC cable directly from the AudioFile.

### Psychophysical procedures

In the psychophysical discrimination experiment, participants heard two amplitude-modulated stimuli presented sequentially using a two-alternative-forced-choice (2AFC) design. The stimulus duration was 500 ms, with a 400 ms interstimulus interval (ISI) and a minimum inter-trial interval of 500 ms. One interval contained the standard stimulus, consisting of the pedestal modulation depth only. The other interval contained the signal stimulus, which comprised the pedestal modulation depth with an additional target increment.

The presentation order of the standard and signal intervals was randomised, and participants were instructed to indicate the interval which they believed contained the target increment using a two-button mouse. A coloured square displayed on the computer screen indicated accuracy (green for correct, red for incorrect). The size of the target increment was determined by a pair of 3-down-1-up staircases, with a step size of 3 dB (where dB units are defined as 20*log_10_(100\**m*)), which terminated after the lesser of 70 trials or 12 reversals. The percentage of correct trials at each target modulation depth was used to fit a cumulative log-Gaussian psychometric function (using Probit analysis) to the data pooled across repetitions. We used this fit to estimate the target modulation that yielded a performance level of 75% correct, which was defined as the threshold. Each participant completed three repetitions of the experiment, producing an average of 223 trials per condition (and an average of 7133 trials in total per participant). This took around 5 hours in total per participant, and was completed across multiple days in blocks lasting around 10 minutes each.

Four binaural arrangements of target and pedestal were tested, at 8 pedestal modulation depths (100\**m* = 0, 1, 2, 4, 8, 16, 32 & 64). The arrangements are illustrated schematically in Figure 2a, and were interleaved within a block at a single pedestal level, so that on each trial participants were not aware of the condition being tested. Note that in all conditions the carrier was presented to both ears, whether or not it was modulated by the pedestal and/or target. This avoids confounding the ears presented with the modulator with those presented with the carrier. In the monaural condition, the pedestal and target modulations were presented to one ear, with the other ear receiving only the unmodulated carrier. The modulated stimulus was assigned randomly to an ear on each trial. In the binaural condition, the pedestal and target modulations were presented to both ears (in phase). Comparison of the binaural and monaural conditions reveals the advantage of stimulating both ears with an AM stimulus, rather than only one. In the dichotic condition, the pedestal modulation was presented to one ear and the target modulation to the other ear. This allows us the measurement of masking effects across the ears. Finally, in the half-binaural condition, the pedestal modulation was played to both ears, but the target modulation to only one ear. When compared with the binaural condition, this arrangement keeps the number of ears receiving the pedestal fixed, and changes only the number of ears receiving the target modulation. It therefore does not confound the effects of pedestal and target stimulation across the ears, and offers a more appropriate comparison than does the monaural condition. Note that for pedestal modulation depths of *m* = 0, and in the dichotic condition, the target increment was relative to the unmodulated carrier. Because the *m* = 0 detection condition was identical across the monaural, dichotic and half-binaural conditions, we pooled data across these conditions to obtain a more reliable estimate of threshold. In all conditions, the modulation frequency for the pedestal and the target was 40 Hz.

### EEG procedure

In the EEG experiment, participants heard 11-s sequences of amplitude-modulated stimuli interspersed with silent periods of 3 seconds. There were five signal modulation depths (*m* = 6.25, 12.5, 25, 50 & 100%) and six binaural conditions, as illustrated in Figure 2b. In the first three conditions, a single modulation frequency (40 Hz, F1) was used. In the monaural condition, the modulated ‘signal’ tone was presented to one ear, and the unmodulated carrier was presented to the other ear. In the binaural condition, the signal modulation was presented to both ears. In the dichotic condition, the signal modulation was presented to one ear, and a modulated masker with a modulation depth of *m* = 50% was presented to the other ear. These three conditions permit estimation of summation and gain control properties, as the use of the same modulation frequency in both ears means that signals to the left and right ears will sum.

The remaining three conditions involved modulation at a second modulation frequency (35 Hz, F2), in order to isolate suppressive processes between channels. In the cross-monaural condition, F2 was presented to one ear as the signal, and the unmodulated carrier was presented to the other ear (F1 was not presented to either ear). This provides a comparison with the 40 Hz monaural condition, and also a baseline with which to compare the other cross-frequency conditions. In the cross-binaural condition, F1 was presented to one ear and F2 was presented to the other ear but the modulation depth of F1 and F2 was the same. This allows measurement of suppressive interactions between the ears without the complicating factor of signal summation at the same modulator frequency tag. In the cross-dichotic condition, F1 was presented to one ear, and F2 (*m* = 50%) was presented to the other ear. Again, we expect this condition to reveal suppressive interactions between the ears, as the F2 mask should suppress the F1 target, and reduce the amplitude of the response measured at 40 Hz.

The order of conditions was randomised, and each condition was repeated ten times, counterbalancing the presentation of stimuli to the left and right ears as required. Trials were split across 5 blocks, each lasting 14 minutes, with rest breaks between blocks. EEG data for each trial at each electrode were then analysed offline. The first 1000 ms following stimulus presentation was discarded to eliminate onset transients, and the remaining ten seconds were Fourier transformed and averaged coherently (taking into account the phase angle) across repetitions. This coherent averaging procedure minimises noise contributions (which have random phase across repetitions), and previous studies [e.g. 35] have indicated that this renders artifact rejection procedures unnecessary. The dependent variables were the signal-to-noise ratios (SNR) at the Fourier components corresponding to the two modulation frequencies used in the experiment (40 Hz, F1 and 35 Hz, F2). These were calculated by dividing the amplitude at the frequency of interest (35 or 40 Hz) by the average amplitude in the surrounding 10 bins (±0.5 Hz in steps of 0.1 Hz). The absolute SNRs (discarding phase information) were then used to average across participants.

### Participants

Six adult participants (two male; age range 22 – 40) completed the psychophysics experiment, and twelve adult participants (3 male; age range 20 – 33) completed the EEG experiment. All had self-reported normal hearing, and provided written informed consent. Experimental procedures were approved by the ethics committee of the Department of Psychology, University of York.

### Data and code sharing

Data and analysis scripts are available online at: https://dx.doi.org/10.17605/OSF.IO/KV2TM

## Results

### Discrimination results are consistent with weak interaural suppression

The results of the AM depth discrimination experiment are shown in Figure 3 averaged across 6 participants. A 4 (condition) x 8 (pedestal level) repeated measures ANOVA found significant main effects of condition (F=47.46, *p*<0.01, η_G_^2^=0.32) and pedestal level (F=10.77, *p*<0.01, η_G_^2^=0.58), and a significant interaction between the two factors (F=8.64, *p*<0.01, η_G_^2^=0.34). When the results are plotted as thresholds on logarithmic axes, the results for binaurally presented modulations (red squares in Figure 3a) followed a ‘dipper’ shape [39], with thresholds decreasing from an average of around 6% at detection threshold to around 2% on a pedestal of 8% (a facilitation effect). At higher pedestal modulations, thresholds increased to more than 16%, indicating a masking effect. Thresholds for the monaural modulation (blue circles in Figure 3a) followed a similar pattern, but were shifted vertically by an average factor of 1.90 across all pedestal levels. The monaural and binaural dipper handles remained apart, and were approximately parallel, at higher pedestal modulation depths. At detection threshold (pedestal *m* = 0), the average summation between binaural and monaural modulation (e.g. the vertical offset between the leftmost points in Figure 3a) was a factor of 1.67 (4.47 dB). This level of summation is above that typically expected from probabilistic combination of independent inputs [40], and implies the presence of physiological summation between the ears.

**Figure 3:**
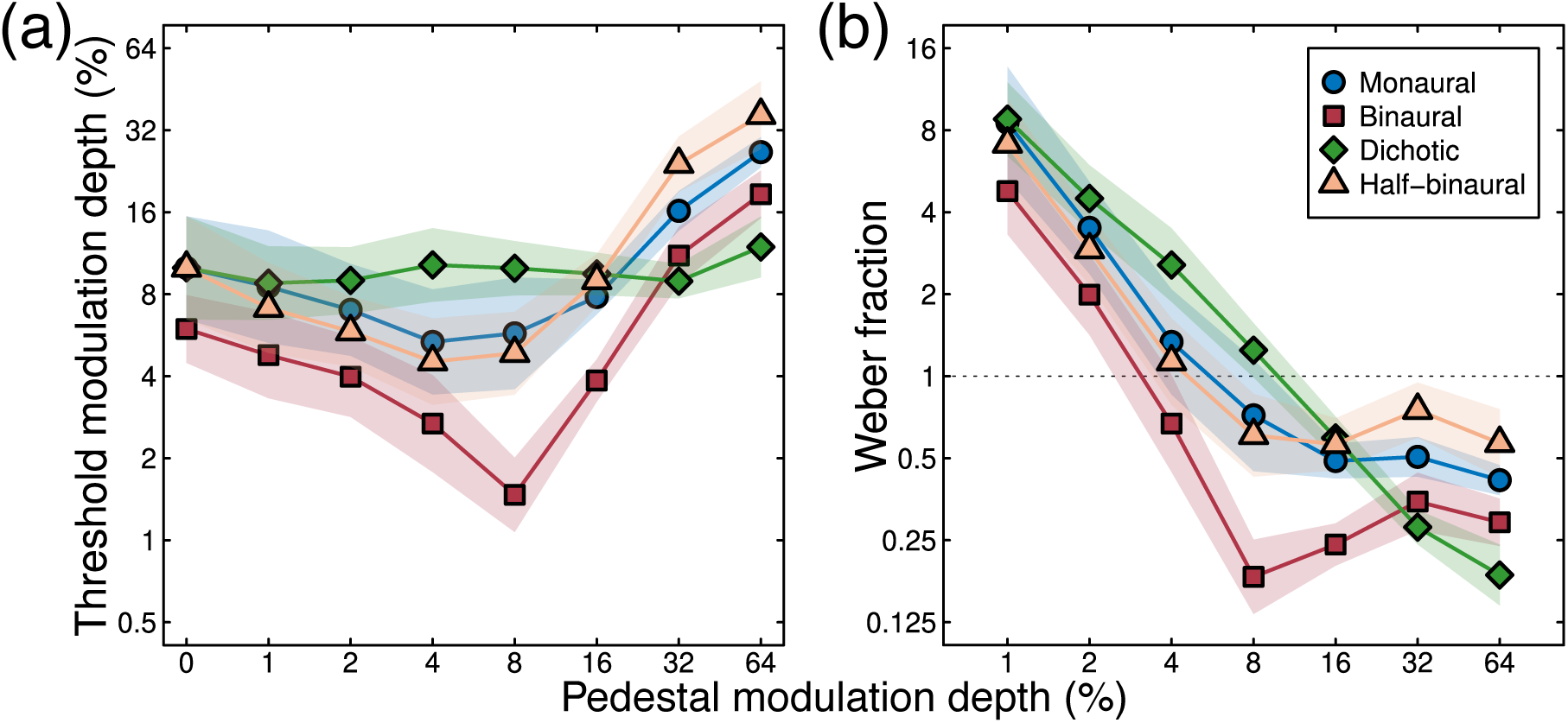
Results of the psychophysical AM depth discrimination experiment averaged across six participants (panel a). Arrangements of pedestal and target in different conditions were as illustrated in Figure 2a. The data are replotted as Weber fractions in panel (b) by dividing each threshold by its accompanying pedestal modulation depth. Shaded regions in both panels give ±1 Standard Error (SE) across participants.

Dichotic presentation (pedestal modulation in one ear and target modulation in the other) elevated thresholds very slightly, by a factor of 1.19 at the highest pedestal modulation depths (green diamonds in Figure 3a), compared to baseline (0% pedestal modulation). This masking effect was substantially weaker than is typically observed for dichoptic pedestal masking in vision (see Figure 1a), which can elevate thresholds by around a factor of 30 [34]. The thresholds for the half-binaural condition (orange triangles in Figure 3a-d), where the pedestal was presented to both ears, but the target only to one ear, was not appreciably different from that for the monaural condition, with thresholds greater than in the binaural condition by a factor of 1.94 on average.

These results can be converted to Weber fractions by dividing the threshold increments by the pedestal modulation depths, for pedestals >0%. These values are shown for the average data in Figure 3b. At lower pedestal modulation depths (<8%), Weber fractions decreased with increasing pedestal level. At pedestal modulations above 8%, the binaural Weber fractions (red squares) plateaued at around 0.25, whereas the monaural and half-binaural Weber fractions (blue circles and orange triangles) plateaued around 0.5. The dichotic Weber fractions (green diamonds) continued to decrease throughout. Thus, non-Weber behaviour occurred over the lower range of pedestal modulations depths, but more traditional Weber-like behaviour was evident at higher pedestal levels. The exception is the dichotic condition, where non-Weber behaviour was evident throughout.

Overall, this pattern of results is consistent with a weak level of interaural suppression between the left and right ears. This accounts for the lack of convergence of monaural and binaural dipper functions at high pedestal levels, and the relatively minimal threshold elevation in the dichotic masking condition, as we will show in greater detail through computational modelling below. Our second experiment sought to measure modulation response functions directly using steady-state EEG to test whether this weak suppression is also evident in cortical responses.

### Direct neural measures of binaural combination

Steady-state EEG signals were evident over central regions of the scalp, at both modulation frequencies tested, and for both monaural and binaural modulations (Figure 4). In particular, there was no evidence of laterality effects for monaural presentation to one or other ear. We therefore averaged steady-state SNRs across a region-of-interest (ROI) comprising nine fronto-central electrodes (Fz, F1, F2, FCz, FC1, FC2, Cz, C1, C2, highlighted white in Figure 4) to calculate modulation response functions.

**Figure 4:**
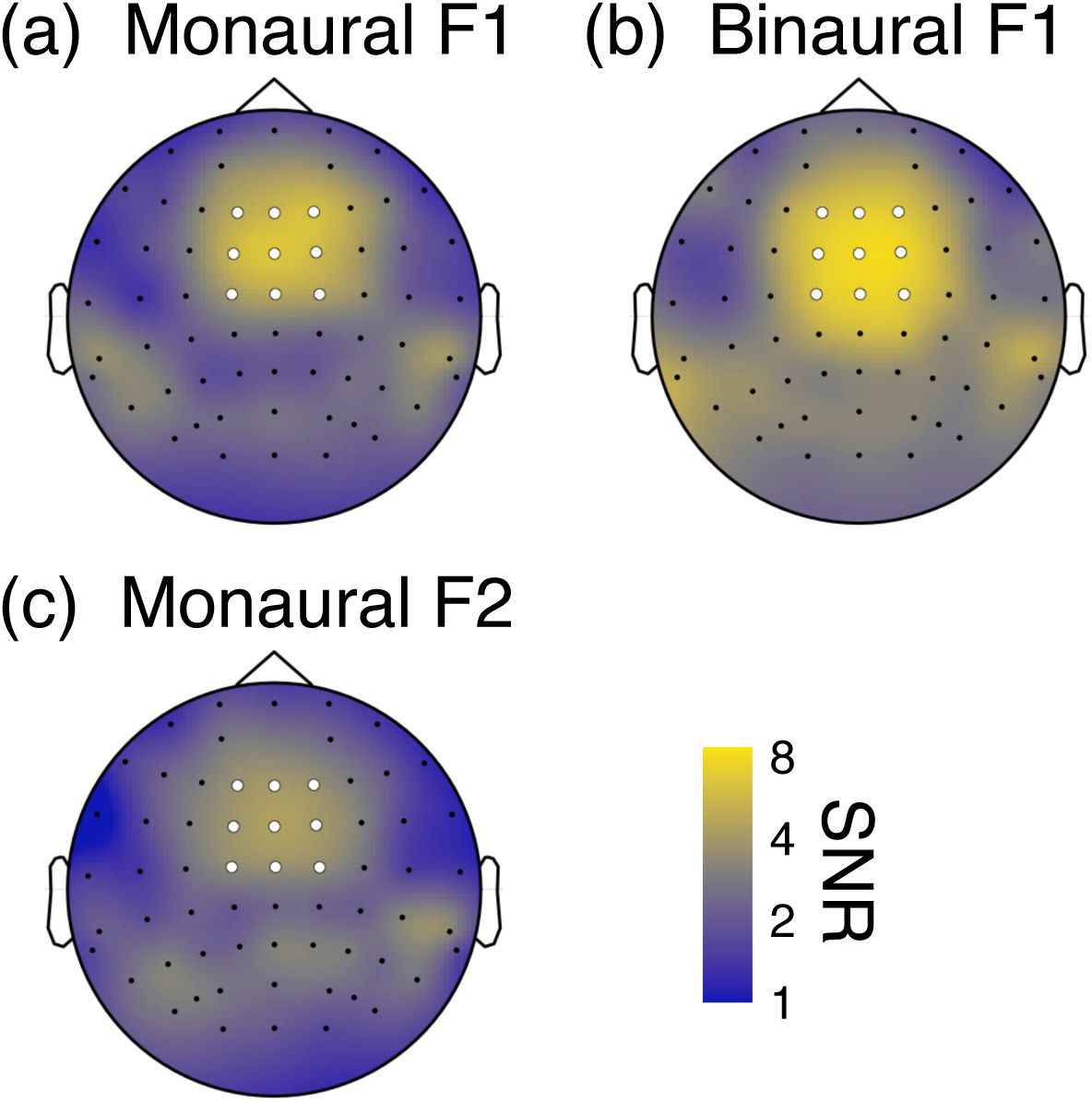
SNRs across the scalp at either 40 Hz (F1, panel a,b) or 35 Hz (F2, panel c). In panels a,c the signal modulation was presented to one ear (averaged across left and right), in panel b the modulation was presented to both ears. Note the log-scaling of the colour map. Dots indicate electrode locations, with those filled white showing the 9 electrodes that comprised the region-of-interest (ROI) used for subsequent analyses.

We conducted separate 6 (condition) x 5 (modulation depth) repeated measures ANOVAs at each modulation frequency using the SNRs averaged across the ROI. At 40 Hz, we found significant main effects of condition (F=38.83, *p*<0.001, η_G_^2^=0.54, Greenhouse-Geisser corrected) and modulation depth (F=33.22, *p*<0.001, η_G_^2^=0.43, Greenhouse-Geisser corrected), and a significant interaction between the two variables (F=6.13, *p*<0.001, η_G_^2^=0.19). At 35 Hz, we also found significant main effects of condition (F=17.40, *p*<0.01, η_G_^2^=0.45, Greenhouse-Geisser corrected) and modulation depth (F=8.72, *p*<0.001, η_G_^2^=0.07), and a significant interaction between the two variables (F=5.64, *p*<0.001, η_G_^2^=0.17, Greenhouse-Geisser corrected).

SNRs are plotted as a function of modulation depth in Figure 5. For a single modulation frequency (40 Hz), responses increased monotonically with increasing modulation depth, with SNRs >2 evident for modulation depths above 12.5%. Binaural presentation (red squares in Figure 5a) led to SNRs of around 7 at the highest modulation depth, whereas monaural modulation produced weaker signals of SNR∼5 (blue circles in Figure 5a). Assuming a baseline of SNR=1 in the absence of any signal (where activity at the signal frequency will equal the noise level in the adjacent frequency bins), this represents a binaural increase in response of a factor of 1.5. The finding that monaural and binaural functions do not converge at high modulation depths suggests that interaural suppression is too weak to fully normalise the response to two inputs compared with one.

**Figure 5:**
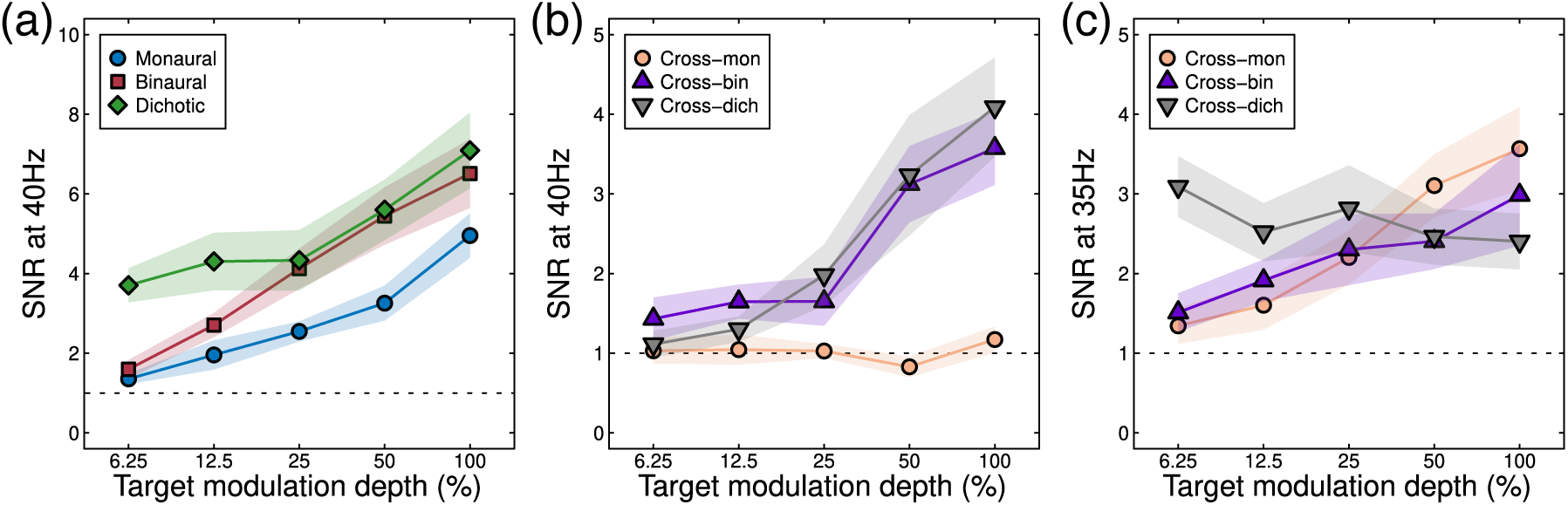
SNRs expressed as a function of signal modulation depth for six conditions at two frequencies. Shaded regions give ±1SE of the mean across participants (N=12). The dashed horizontal line in each plot indicates the nominal baseline of SNR=1.

In the dichotic condition (green diamonds in Figure 5a), a masker with a fixed 50% modulation depth presented to one ear produced an SNR of 4 when the unmodulated carrier was presented to the other ear (see left-most point). As the dichotic signal modulation increased, responses increased to match the binaural condition at higher signal modulations (red squares and green diamonds converge in Figure 5a).

When the carrier presented to one ear was modulated at a different frequency (35 Hz), several differences were apparent for the three conditions. Monaural modulation at 35 Hz (the cross-mon condition) evoked no measureable responses at 40 Hz as expected (orange circles in Figure 5b). At the modulation frequency of 35 Hz, this condition produced a monotonically increasing function peaking around SNR=3.5 (orange circles in Figure 5c). Binaural modulation with different modulation frequencies in each ear led to weaker responses (SNRs of 4 at 40 Hz and 3 at 35 Hz; purple triangles in Figure 5b,c) than for binaural modulation at the same frequency (SNR=7, red squares in Figure 5a). A 35 Hz AM masker with a fixed 50% modulation depth presented to one ear produced little change in the response to a signal in the other ear, which was amplitude-modulated with a modulation frequency of 40 Hz (grey inverted triangles in Figure 5b), though increasing the signal modulation depth slightly reduced the neural response to the 35-Hz AM masker (grey inverted triangles in Figure 5c). This weak dichotic masking effect is further evidence of weak interaural suppression. We next consider model arrangements that are able to explain these results.

### A single model of signal combination predicts psychophysics and EEG results

To further understand our results, we fit the model described by equation 1 to both data sets. To fit the psychophysical data, we calculated the target modulation depth that was necessary to increase the model response by a fixed value, *σ*_40_, which was a fifth free parameter in the model (the other four free parameters being *p, q, Z* and *ω*; note that all parameters were constrained to be positive, *q* was constrained to always be greater than 2 to ensure that the nonlinearity was strong enough to produce a dip, and we ensured that *p*>*q*). With five free parameters, the data were described extremely well (see Figure 6a), with a root mean square error (RMSE, calculated as the square root of the mean squared error between model and data points across all conditions displayed in a figure panel) of 1.2 dB, which compares favourably to equivalent model fits in vision experiments [34]. However, the value of the interaural suppression parameter was much less than 1 (*ω* = 0.02, see Table 1). This weak interaural suppression changes the behaviour of the canonical model shown in Figure 1c in two important ways, both of which are consistent with our empirical results. First, the degree of threshold elevation in the dichotic condition is much weaker, as is clear in the data (green diamonds in Figures 3a & 6a). Second, the thresholds in the monaural condition are consistently higher than those in the binaural condition, even at high pedestal levels (compare blue circles and red squares in Figures 3a & 6a).

**Table 1:** Parameters for the model fits shown in Figure 6 with parameter constraints as described in the text.

| Panel | $p$ | $q$ | $Z$ | $\sigma_{40}$ | $\sigma_{35}$ | $\omega$ | RMSE |
| --- | --- | --- | --- | --- | --- | --- | --- |
| 6a | 2.86 | 2.47 | 10.22 | 0.88 | - | 0.02 | 1.2dB |
| 6b | 2.86 | 2.47 | 10.22 | 0.88 | - | 1 | 5.5dB |
| 6c | 2.13 | 2.00 | 25.16 | 0.15 | - | 1 | 4.3dB |
| 6d | 2.41 | 2.00 | 10.94 | 1.85 | 2.55 | 0.14 | 0.34 |
| 6e | 2.41 | 2.00 | 10.94 | 1.85 | 2.55 | 1 | 0.96 |
| 6f | 2.00 | 2.00 | 47.26 | 0.18 | 0.28 | 1 | 0.51 |

**Figure 6:**
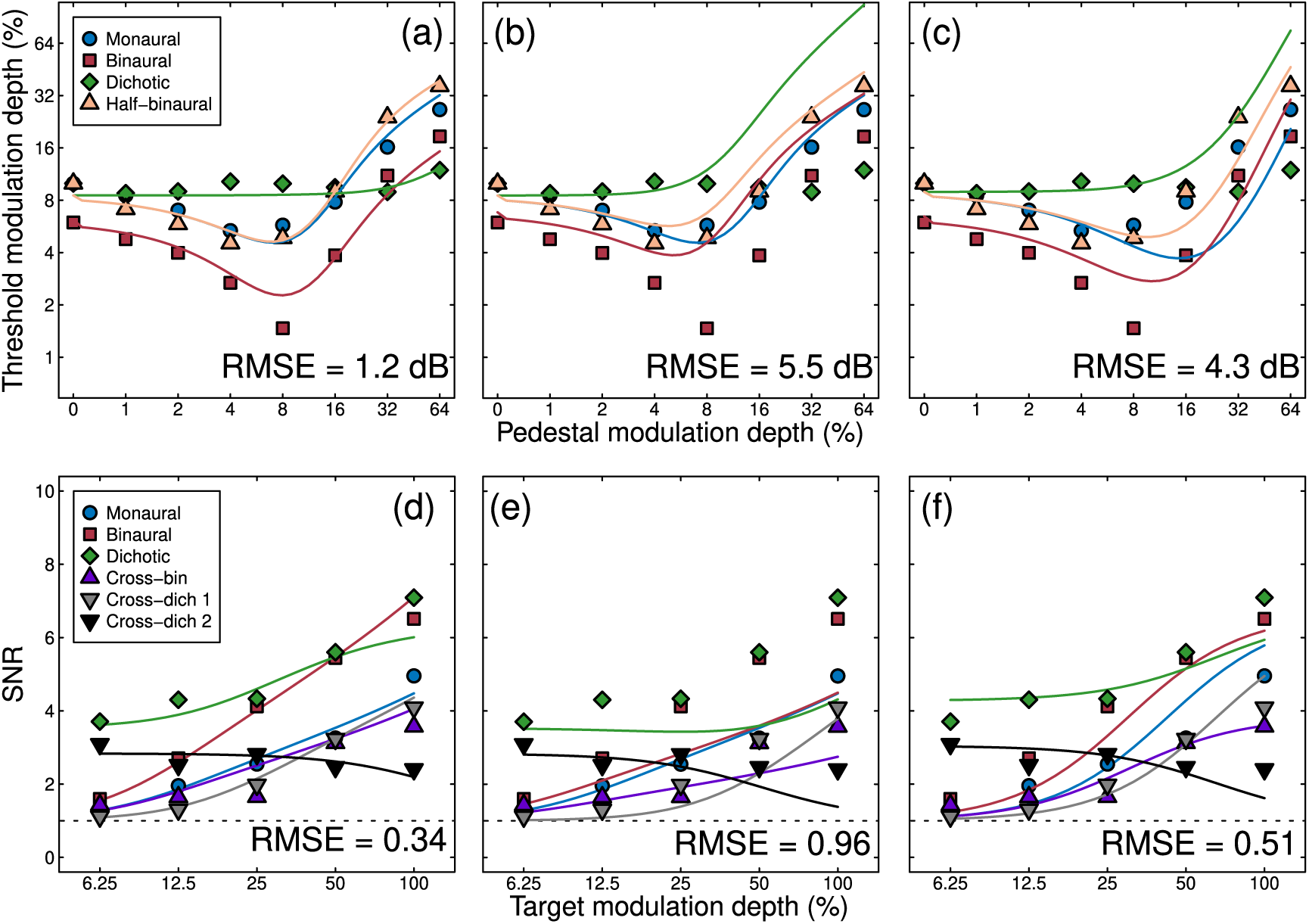
Fits of several variants of the signal combination model (curves) to empirical data (symbols). Model fits in panels (a,d) had all parameters free to vary. Model curves in panels (b,d) retained the fitted parameters from (a,d), but altered the weight of interaural suppression to *ω* = 1. Model fits in panels (c,f) had all parameters free to vary apart from the weight of interaural suppression, which was fixed at *ω* = 1. The data in the top panels are the averaged dipper functions duplicated from panel 3a, and those in the lower row are collapsed across the three panels of Figure 5 (omitting the mon-cross condition). Values in the lower right of each plot give the root mean square error (RMSE) of the fit in logarithmic (dB) units in the upper panels, and in units of SNR in the lower panels.

To illustrate how the model behaves with stronger interaural suppression, we increased the weight to a value of *ω* = 1, but left the other parameters fixed at the values from the previous fit. This manipulation (shown in Figure 6b) reversed the changes caused by the weaker suppression – masking became stronger in the dichotic condition, and the monaural and binaural dipper functions converged at the higher pedestal levels. These changes provided a poorer description of human discrimination performance, with the RMSE increasing from 1.2 dB to 5.5 dB. Finally, we held suppression constant (at *ω* = 1), but permitted the other four parameters to vary in the fit. This somewhat improved the fit (see Figure 6c), but retained the qualitative shortcomings associated with strong interaural suppression, and only slightly improved the RMSE (from 5.5 dB to 4.3 dB).

To fit the EEG data, we converted the model response to an SNR by adding the noise parameter (*σ*) to the model response, and then scaling by the noise parameter (e.g. (resp + *σ*)/*σ*). Because maximum SNRs varied slightly across the two modulation frequencies (40 and 35 Hz, see Figure 5), we permitted this noise parameter to take a different value at each frequency (*σ*_40_ and *σ*_35_). Model predictions for the conditions described in Figure 2b are shown in Figure 6d for a version of the model with six free parameters. This produced an excellent fit [comparable to those for visual signals, see 35], which included the main qualitative features of the empirical amplitude response functions, with an RMSE of 0.34. The model captures the increased response to binaural modulations compared with monaural modulations (blue circles vs red squares in Figure 6d), the relatively modest suppression in the cross-bin (purple triangles) and cross-dichotic (grey triangles) conditions at 40 Hz relative to the monaural condition, and the gentle decline in SNR in the cross dichotic condition at the masker frequency (black triangles in Figure 6d). Most parameters took on comparable values to those for the dipper function fits described above (see Table 1). Of particular note, the weight of interaural suppression remained weak (*ω* = 0.14).

We again explored the effect of increasing the weight of suppression (to *ω* = 1) whilst keeping the other parameters unchanged. This resulted in a reduction of amplitudes in the binaural and cross-binaural conditions, which worsened the fit (to an RMSE of 0.96). Permitting all other parameters (apart from *ω*) to vary freely improved the fit (to RMSE = 0.51), but there were still numerous shortcomings. In particular the monaural and binaural response functions were more similar than in the data, and the reduction in SNR in the cross-binaural and cross-dichotic conditions was more extensive than found empirically.

Our modelling of the data from two experimental paradigms therefore support the empirical finding that interaural suppression is relatively weak (by around an order of magnitude) compared with analogous phenomena in vision (interocular suppression).

## Discussion

We have presented converging evidence from two experimental paradigms (psychophysics and steady-state EEG) concerning the architecture of the human binaural auditory system. A single computational model, in which signals from the two ears inhibit each other weakly before being combined, provided the best description of data sets from both experiments. This model architecture originates from work on binocular vision, showing a commonality between these two sensory systems. We now discuss these results in the context of related empirical results, previous binaural models, and ecological constraints that differentially affect vision and hearing.

### A unified framework for understanding binaural processing

Our psychophysical experiment replicates the classical finding [1,6–9] of approximately 3 dB of binaural summation at detection threshold (here 4.47 dB) for amplitude-modulated stimuli. This is very similar to values previously reported for binocular summation of contrast in the visual system, where summation ratios of 3 – 6 dB are typical [41]. Above threshold, this difference persisted, with monaural stimulation producing higher discrimination thresholds (Figure 3) and weaker EEG responses (Figures 4, 5) than binaural stimulation. This is consistent with previous EEG work [17,22], and also the finding that perceived loudness and modulation depth are higher for binaural than monaural presentation [11–14,27]. However, these auditory effects are dramatically different from the visual domain, where both discrimination performance and perceived contrast are largely independent of the number of eyes stimulated [33]. We discuss possible reasons for this modality difference below.

Suppression between the ears has been measured previously with steady-state magnetoencephalography (MEG) using amplitude-modulated stimuli with frequencies that are the same [42] or different [23,24] in the left and right ears. When the same frequency is used in both ears, suppression can be assessed by comparing binaural responses to the linear sum of two monaural responses. When the measured binaural response is weaker than this prediction, this is taken as evidence of suppression between the ears (though we note that nonlinear transduction might produce similar effects). Tiihonen et al. [42] used 500 ms click trains at 40 Hz, and found evidence for strong suppression of the initial evoked N100 amplitudes, but weaker suppression of the 40 Hz response (especially relative to ipsilateral stimuli). If suppression decreased even further for longer presentations (as used here), this might explain why suppression appears so weak in our study. Alternatively, the Tiihonen study used laterally placed MEG sensors to record signals from auditory cortex, whereas we used EEG with a central region of interest, which might also account for the differences. Two other studies [23,24] used different frequency tags in the two ears in conditions analogous to our cross-binaural and cross-dichaural conditions. For tag frequencies around 20 Hz, there were varying amounts of suppression between 36% and 72% of the monaural response depending on whether signals were measured from the left or right hemisphere, and whether they were for ipsilateral or contralateral presentations [23]. A second study [24] used frequencies around 40 Hz, and again found a range of suppression strengths depending on laterality and hemisphere. The weakest suppressive effects were comparable to those measured here using steady-state EEG (see Figure 5b,c). It is possible that different stages of processing might involve different amounts of suppression, which would require the use of techniques with better spatial precision to localise responses of interest to specific brain regions.

Another widely-studied phenomenon that might involve suppression between the ears is the binaural masking level difference [BMLD; 43]. In this paradigm, a signal embedded in noise is detected more easily when either the signal or the noise in one ear is inverted in phase [44]. Contemporary explanations of this effect [45,46] invoke cross-correlation of binaural signals, but lack explicit inhibition between masker and test signals. However, more elaborate versions of the model described here include mechanisms tuned to opposite spatial phases of sine-wave grating stimuli [47], and a similar approach in the temporal domain might be capable of predicting BMLD effects. Alternatively, since the BMLD phenomenon often involves segmentation of target and masker, it might be more akin to ‘unmasking’ effects that occur in vision when stimuli are presented in different depth planes [48,49]. Modelling such effects would likely require additional mechanisms representing different spatial locations, far beyond the scope of the architecture proposed here.

### The model shares features with previous binaural models

Previous models of binaural processing [50–52] have some architectural similarities to the model shown in Figure 1a. For example, binaural inhibition is a common feature [52], often occurring across multiple timescales [50]. However these models are typically designed with a focus on explaining perception across a range of frequencies (and for inputs of arbitrary frequency content), rather than attempting to understand performance on specific tasks (i.e. AM depth discrimination) or the precise mapping between stimulus and cortical response (i.e. the amplitude response functions measured using steady-state EEG). At threshold, one model [51] predicts minimal levels of binaural summation (∼1dB) in line with probabilistic combination of inputs but below that found experimentally. These models would therefore likely require modification (i.e. the inclusion of physiological summation and early nonlinearities) to explain the data here, though it is possible that such modifications could be successful, given the other similarities between the models.

Several previous neural models of binaural processing have focussed on excitatory and inhibitory processes of neurons in subcortical auditory structures such as the lateral superior olive. These models (reviewed in [45]) are concerned with lateralised processing, in which interaural interactions are purely inhibitory, and so do not typically feature excitatory summation. However, models of inferior colliculus neurons do typically involve binaural summation, and have the same basic structure as the architecture shown in Figure 1a. In general these models are designed to explain responses to diotically asynchronous stimuli (where stimuli reach the two ears at different times), and so typically feature asymmetric delays across the excitatory and inhibitory inputs from the two ears [e.g. 53]. Since a time delay is not a critical component of the divisive suppression on the denominator of equation 1, and because a mechanism with broad temporal tuning is equivalent to the envelope of many mechanisms with different delays, the architecture proposed here can be considered a generalised case of such models.

### Ecological constraints on vision and hearing

This study reveals an important and striking difference between hearing and vision – suppression between the ears is far weaker than suppression between the eyes. Why should this be so? In the visual domain, the brain attempts to construct a unitary percept of the visual environment from two overlapping inputs, termed binocular single vision. For weak signals (at detection threshold) it is beneficial to sum the two inputs to improve the signal-to-noise ratio. But above threshold, there is no advantage for a visual object to appear more intense when viewed with two eyes compared with one. The strong interocular suppression prevents this from occurring by normalizing the signals from the left and right eyes to achieve ‘ocularity invariance’ – the constancy of perception through one or both eyes [33]. The guiding principle here may be that the brain is reducing redundancy in the sensory representation by avoiding multiple representations of a single object.

In the human auditory system the ears are placed laterally, maximising the disparity between the signals received (and minimising overlap). This incurs benefits when determining the location of lateralised sound sources, though reporting the location of pure tone sources at the midline (i.e. directly in front or behind) is very poor [2]. Hearing a sound through both ears at once therefore does not necessarily provide information that it comes from a single object, and so the principle of invariance should not be applied (and interaural suppression should be weak). However other cues that are consistent with a single auditory object (for example interaural time and level differences consistent with a common location) should result in strong suppression to reduce redundant representations, and cues that signals come from multiple auditory objects should release that suppression. This is the essence of the BMLD effects discussed above – suppression is strongest when target and masker have the same phase offsets (consistent with a common source), and weakest when their phase offsets are different. The distinct constraints placed on the visual and auditory systems therefore result in different requirements, which are implemented in a common architecture by changing the weight of suppression between channels.

## Conclusions

A combination of psychophysical and electrophysiological experiments, and computational modelling have converged on an architecture for the binaural combination of amplitude-modulated tones. This architecture is identical to the way that visual signals are combined across the eyes, with the exception that the weight of suppression between the ears is weaker than that between the eyes. This is likely because the ecological constraints governing suppression of multiple sources aim to avoid signals from a common source being over-represented. Such a high level of consistency across sensory modalities is unusual, and illustrates how the brain can adapt generic neural circuits to meet the demands of a specific situation.

## Acknowledgements

This work was supported by the Royal Society (grant number RG130121 to DHB) and the Wellcome Trust (grant number 213616/Z/18/Z to AB). This work was also supported by the NIHR Manchester Biomedical Research Centre.

## Supplementary Materials

**Figure S1:**
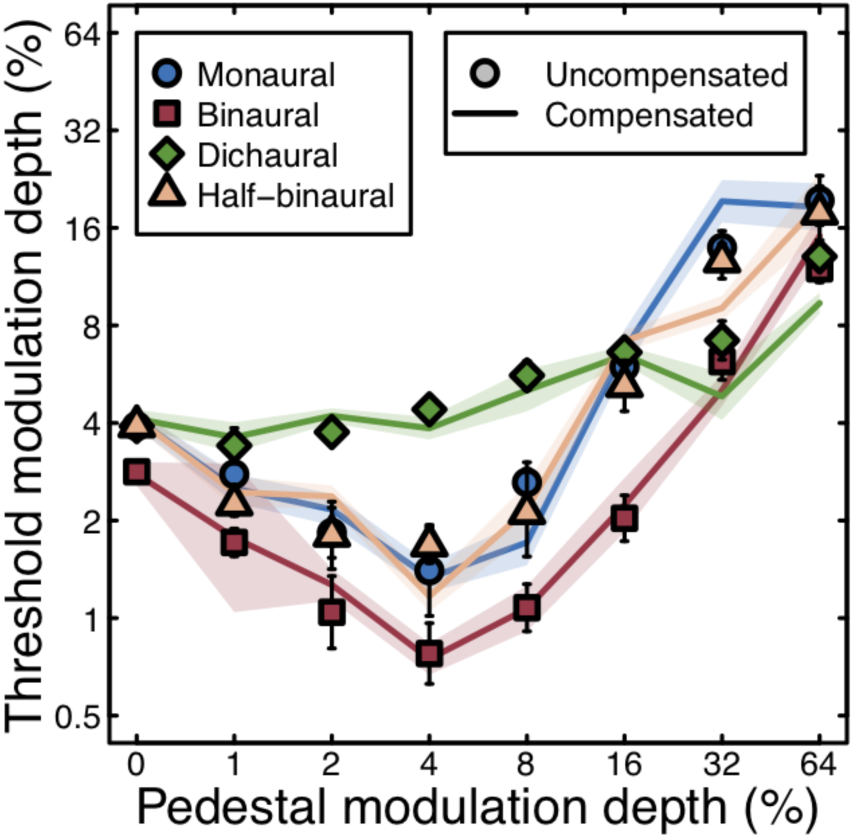
Comparison of thresholds for uncompensated (symbols) and compensated (lines) amplitude modulated stimuli, for one participant. We compensated for overall stimulus power using the method of Ewert and Dau [37], in which each stimulus was scaled by 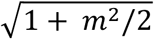, where *m* is the modulation depth. Overall, the compensation had no systematic effect on thresholds. A paired t-test across all conditions produced no significant difference (*t*=0.26, *df*=31, *p*=0.79). Error bars and shaded regions give ±1SE of the Probit fit.

